# Diversity and prevalence of *Clostridium innocuum* in the human gut microbiota

**DOI:** 10.1101/2022.06.29.498201

**Authors:** Disha Bhattacharjee, Clara Flores, Christine Woelfel-Monsivais, Anna M. Seekatz

**Author notes:** Corresponding Author. Life Sciences Building 157A, 190 Collings St, Clemson, South Carolina – 29634, United States of America.

## Abstract

Clostridia are a polyphyletic group of Gram-positive, spore-forming anaerobes in the Firmicutes phylum that significantly impact metabolism and functioning of human gastrointestinal tract. Recently, Clostridia were divided into two separate classes, Clostridia and Erysipelotrichia, based on phenotypic and 16S rRNA gene-based differences. While Clostridia include many well-known pathogenic bacteria, Erysipelotrichia remain relatively uncharacterized, particularly regarding their role as a pathogen vs. commensal. Despite wide recognition as a commensal, the Erysipelotrichial species, *Clostridium innocuum*, has recently been associated with various disease states. To further understand the ecological and potential virulent role of *C. innocuum*, we conducted a genomic comparison across 38 *C. innocuum* isolates and 75 publicly available genomes. Based on colony morphology, we isolated multiple *C. innocuum* cultivars from the feces of healthy human volunteers (n=5). Comparison of the 16S rRNA gene of our isolates against publicly available microbiota datasets in healthy individuals suggests a high prevalence of *C. innocuum* across the human population (> 80%). Analysis of single nucleotide polymorphisms (SNPs) across core genes and average nucleotide identify (ANI) revealed the presence of 4 clades among all available unique genomes (n=108 total). Investigation of carbohydrate and protein utilization pathways, including comparison against the carbohydrate-activating-enzyme (CAZyme) database, demonstrated inter-and intra-clade differences that were further substantiated *in vitro*. Collectively, these data indicate genetic variance within the *C. innocuum* species that may help clarify its role in human disease and health.

**IMPORTANCE:** Clostridia are a group of medically important anaerobes as both commensals and pathogens. Recently, a new class of Erysipelotrichia containing a number of re-assigned Clostridial species has emerged, including *Clostridium innocuum*. Recent studies have implicated *C. innocuum* as a potential causative agent of diarrhea in patients from whom *Clostridioides difficile* could not be isolated. Using genomic and *in vitro* comparison, this study sought to characterize *C. innocuum* in the healthy human gut. Our analyses suggest that *C. innocuum* is a highly prevalent and diverse species, demonstrating clade-specific differences in metabolism and potential virulence. Collectively, this study is the first investigation into a broader description of *C. innocuum* as a human gut inhabitant.

## INTRODUCTION

Commensal bacteria, viruses, fungi, and protozoa, collectively termed microbiota, dominate all surfaces of multicellular hosts, providing them with essential functions. The collective genes provided by individual or groups of microbes maintain health of the host, providing functions such as colonization resistance against pathogens via multiple mechanisms, including nutrient niche exclusion (3–5), modulation of oxygen or pH gradients along the gut (6, 7), or production of metabolites that harm pathogens (8, 9). However, variability of the gut microbiota across individual hosts (10, 11) and lack of characterization of many common gut inhabitants (1) complicate discernible conclusions about many individual members. Recent genomic and phenotypic studies have highlighted strain-level diversity within prominent gut species that can extricate our understanding of the microbiota in health (12–14), which is not captured by 16S rRNA gene-based surveys.

The human gut microbiota is predominantly occupied by anaerobic bacteria, with the most abundant phyla being Firmicutes and Bacteroidetes (15). The diversity and function of several prevalent Bacteroidetes species has been extensively investigated (16), leading to their use as prominent model organisms to understand gut microbiota function (17). For example, species within the *Bacteroides* genus are known to harbor hundreds of Polysaccharide Utilization Loci (PULs) that degrade different glycans and carbohydrates (17), ultimately supplying nutrients to both the host and surrounding microbes that contribute to protection from pathogens. Many prevalent taxa within the polyphyletic, gram-positive Firmicutes phylum remain more nebulous. Within the gut, the Firmicutes phylum is comprised of three classes based on analyses of 16S rRNA nucleotide sequence: Bacilli, Clostridia and Erysipelotrichia (18–20). Bacilli and Clostridia comprise of well-studied members with major implications in industrial applications, health, and disease (2, 21–23). Clostridia as a group have been demonstrated to induce beneficial immune responses, in part via their ability to produce short chain fatty acids that can attenuate gut inflammation (25, 26). In comparison, the importance of Erysipelotrichia in the human gut microbiota remains relatively unexplored. Erysipelotrichia include species that share a genomic resemblance to Mollicutes, a class of parasitic bacteria that are characterized by their distinct lack of cell walls compared to the phylum Tenericutes (27). As a group, Erysipelotrichia in the gut have been associated with host lipid metabolism (28, 29) and disease in humans (30). In mice, expansion of Erysipelotrichia species have been observed following antibiotic treatment (31) or when fed a Western-diet (32).

The role of the Erysipelotrichial species *Clostridium innocuum* in human health remains especially ambiguous. *C. innocuum* was first isolated from an appendiceal abscess but deemed innocuous due to lack of virulence observed in mice and guinea pigs (33). Recently re-classified from its original Clostridial designation (34), *C. innocuum* has been identified as part of the “normal” gut microbiota via its capability to biodegrade glucoseureide (35). Although the first phenotypic description of the organism suggest a non-motile, non-virulent nature of *C. innocuum* (33), current literature suggests otherwise. It has been implicated in extra-intestinal infection and *Clostridioides difficile*-like antibiotic associated diarrhea (35–37), as well as in case studies of bacteremia, endocarditis, osteomyelitis, and peritonitis (28, 38–40). Most recently, a study on Crohn’s disease conducted in mice identified *C. innocuum* in inflamed intestinal tissue of patients with Crohn’s disease (41). Despite these studies, a direct virulence mechanism has yet to be identified (37, 41–43).

Given the putative prevalence of *C. innocuum* in the human gut microbiota, we aimed to investigate the genomic and phenotypic diversity of *C. innocuum*. We compared genomic phylogeny, functional capacity, and virulence factors across single isolates and publicly available genomes. Using a custom 16S rRNA database, we identified *C. innocuum* as a highly prevalent human gut inhabitant. Single nucleotide polymorphisms in core genes suggest that the *C. innocuum* species splits into multiple clades, characterized by differences in metabolism and potential virulence. While comparison to known virulence factors did not identify a direct link to previously associated disease studies, we did identify clade-specific putative virulence factors. Together, these data support a role for *C. innocuum* and related Erysipelotrichial species in modulating the gut nutrient landscape, as well as a strain-specific role for potential virulence.

## MATERIALS AND METHODS

### Isolation of *C. innocuum*

This study was approved by Clemson University’s Institutional Review Board. Healthy donors were over 18, had not taken antibiotics or been diagnosed with any infections within six months, and were not immunocompromised or diagnosed with chronic gastrointestinal conditions. Upon receipt, fecal samples were placed under anaerobic conditions (Coy Laboratory Products, Grass Lake, MI; 85% nitrogen, 10% hydrogen, 5% carbon dioxide) and streaked onto Brain Heart Infusion (BHI) (44), BHI supplemented with fetal bovine serum (FBS; 50 mL/L BHI), or Taurocholate Cycloserine-Cefoxitin-Fructose (TCCFA) (45, 46) using different streaking strategies. Streaks were incubated at 37°C for at least 24 hours, then picked and streaked for purity. Samples were stored at −80°C in 20% glycerol stocks for future *in vitro* work or DNA extraction. Additional details are available in Supplemental Methods.

### DNA extraction and identification of *C. innocuum*

All isolates were heat-extracted at 95°C for 20 minutes for polymerase chain reaction (PCR) using GoTaq (Promega M7132) and 8F and 1492R primers to amplify the whole 16S rRNA gene (4). PCR products were cleaned up using Illustra™ ExoProStar™ kit (Cytiva US78210) and sent to Eton Biosciences for Sanger sequencing, using EzBiocloud and RDP databases for identification (47, 48). DNA extraction for sequencing was performed from 1.8 mL of overnight culture using the Qiagen DNAeasy UltraClean Microbial kit (Qiagen 12224-250). Extracted DNA was diluted to 10 ng/μL concentration (Qubit, Life Technologies, Catalogue #Q33230) and sent to the Microbial Genome Sequencing Center (MiGS), Pittsburgh (www.migscenter.com) for Illumina sequencing using the NextSeq 2000 platform.

### Prevalence of *C. innocuum* in human 16S rRNA gene-based surveys

Fasta sequences of full-length 16S rRNA sequences from *C. innocuum* genomes were formatted for alignment in mothur (49) and aligned using the SILVA database (v132) (50). Previously published sequences from fecal microbiota samples representing ‘healthy’ (51), hospitalized (52), or antibiotic-exposed (53) individuals were processed in mothur using the Schloss lab SOP, aligning to the SILVA database (54), then classifying to the custom classifier using the ‘classify.seqs’ command in mothur (cutoff=95) or directly to the RDP database (v16) for comparison (56). The log_10_ relative abundance of *C. innocuum* was plotted in R using the Kruskal-Wallis test in R with a pairwise Wilcoxon Rank Sum Test for pairwise comparisons between groups.

### *In vitro* growth of *C. innocuum*

Representative strains were chosen from all available *C. innocuum* strains based on 99% dereplication using pyANI in anvi-dereplicate-genomes (57, 58) (Table S2). Strains from freezer stocks were initially streaked onto TCCFA in an anaerobic chamber and incubated at 37°C for 24 hours. A single colony was added into 4 mL TCCF broth (TCCFB) and incubated at 37°C for 24 hours. 1.8 mL of overnight culture was centrifuged at 6,000 rpm for 5 min. 10 μL of the resuspended pellet in 1.8 mL of pre-reduced water strain was added into wells of a 96-well plate (Costar^TM^) containing 100 μl basal media (59) with casamino acids (BMCA) with or without selected carbohydrate sources at 4% w/v (Table S3). A positive control of the resuspended strain in TCCF broth, and BMCA without strain was included on each plate. The prepared plate was placed into a plate reader (Tecan Sunrise^TM^) for growth at 37°C, measuring OD_600_ every 15 minutes for 24 hours. A bromocresol purple (BCP) assay was used on the plate growth to assess pH (Supplemental Methods).

### Whole genome assembly and phylogeny

Full commands are available on https://github.com/SeekatzLab/C.innocuum-diversity, with package versions listed in Table S4, and further description in Supplemental Methods. Briefly, raw reads were quality-checked and adapter-trimmed using Trim-galore (60), assembled using SPAdes (61) as optimized with MEGAHIT (62). Quast with MultiQC was used to calculate assembly statistics (Table S1) (63, 64). Average coverage was calculated using Bowtie2 and SAMtools (65, 66). Prokka was used to annotate assemblies (67). To verify the assembly identity, annotations were run through NCBI Blast and EzBioCloud. Assemblies were also mapped on to the Genome Taxonomy Database (GTDB) (68) through GTDB-tk using Peptostreptococcaceae (i.e., containing *Clostridioides difficile*) as the taxon outgroup (69). Maximum likelihood trees from the *C. innocuum* core genome SNP sites was determined by Roary using RaXML 8.2.12 bootstrapping 500 times. The 16S rRNA maximum likelihood tree was aligned using Clustal Omega and bootstrapped 500 times by RAxML (70, 73). The amino acid fasta phylogenetic tree was mapped against the “phylophlan” database with DIAMOND in Phylophlan (74). Trees were visualized using Graphlan (75) or ggtree and treeio packages in R (76, 77).

### Pangenome analysis, functional enrichment, average nucleotide identity, and dereplication

Contigs from SPAdes were reformatted, and annotated with the COG and KEGG database using Anvi’o version 7.0 (58). Anvi’o was also used to create and visualize the pangenome, determine average nucleotide identity (ANI), and dereplicate strains within the dataset (see Supplemental Methods for details). Heap’s law was calculated in R (formulated as n = κN^γ^, where n is the pan-genome size, N is the number of genomes used, and κ and γ are the fitting parameters) and the α parameter was calculated using micropan (82, 83). Average nucleotide identity (ANI) was computed using the anvi-compute-genome-similarity with pyANI (57). Dereplication between strains was computed using anvi-dereplicate-genomes at 90, 95, 98, 99, 99.9 and 100 % similarity threshold (Table S2). Ten strains from the entire set depreplicated at 100%, so only one of those copies were kept for a total of 108 unique genomes. False Detection Rate correction to p-values for functional enrichment was applied using the package qvalue from Bioconductor (78) and visualized using dplyr, ggplot2, and readxl packages (79–81).

### CAZyme and putative virulence

CAZymes were predicted using DBCAN version 2.0.6 (84), using the Fasta nucleotide sequences generated from Prokka for each of the strains. Prokka generated nucleotide fasta files (.fna) were processed through PathoFact (85) to predict virulence factors, toxins, and antimicrobial peptides. Genomic islands were predicted in *C. innocuum* 14501 using the web computational tool IslandViewer 4 (86). Circos was used to visualize chromosomal locations (86).

### Data Availability

All raw sequence data and associated information has been deposited in the NCBI Sequence Read Archive under BioProject PRJNA841489. All code used to analyze data are available at https://github.com/SeekatzLab/C.innocuum-diversity.

## RESULTS

### *Clostridium innocuum* is a prevalent human gut microbe

We screened five fresh fecal samples for the presence of *C. innocuum* strains as part of a larger study focused on cultivation of gut isolates (Figure 1A). Sanger sequencing of the full-length 16S rRNA gene from morphologically distinct colonies confirmed the presence of 38 isolates that matched *C. innocuum* (80% similarity) (Table S1), with each fecal sample yielding at least three distinct colonies. While metagenomic and 16S rRNA gene-based surveys suggest *C. innocuum* may be a common resident of the human gut microbiota, its prevalence across the human population is unknown. To broadly identify the presence of *C. innocuum* within the human gut microbiota, we compared multiple available 16S rRNA datasets from previous human gut microbiota studies to a curated database consisting of the 16S gene from our isolates (49, 54). Within these datasets, approximately 80% of samples (n = 420) contained *C. innocuum* sequences, suggesting a higher prevalence of *C. innocuum* within the healthy human gut microbiota than previously appreciated (Figure 1B). Although the relative abundance (RA) of *C. innocuum* in feces retrieved from 16S rRNA gene-based sequencing data was relatively low across all samples (mean RA = 0.22%), samples collected from patients on antibiotics (mean RA = 0.40%; n = 96) or with sepsis (mean RA = 0.46%; n = 24) were significantly increased compared to healthy controls (mean RA = 0.16%; n = 250).

**Figure 1.**
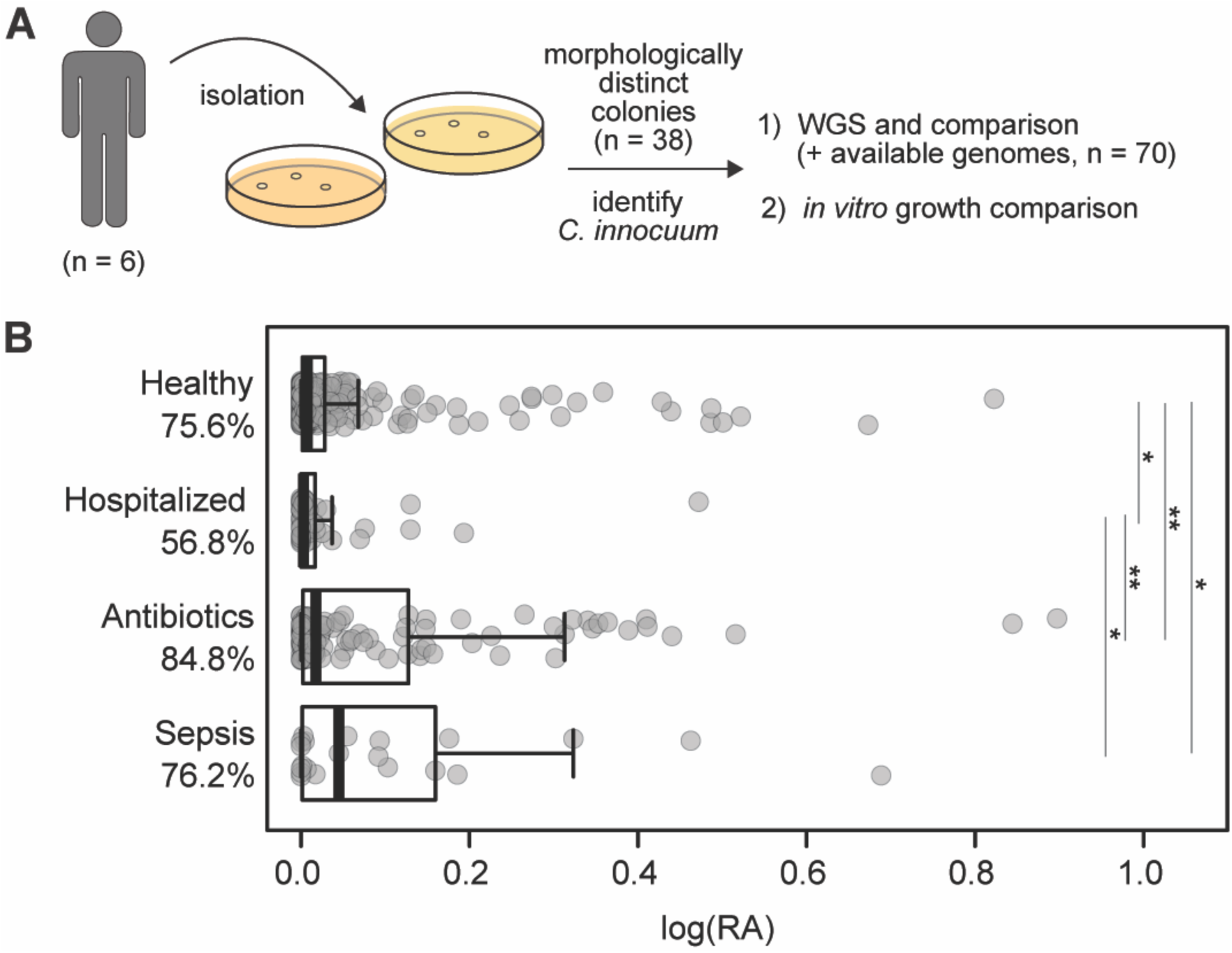
*C. innocuum* is a prevalent human gut bacterium. **A)** Isolation pipeline design for *C. innocuum*. **B)** Detection of *C. innocuum* in human feces across previously published 16S rRNA datasets using a custom classifier, categorized by published disease status. Log_10_ of relative abundance is displayed on the x-axis, with prevalence (% detected based on presence/absence). Pairwise Wilcoxon Rank Sum Test, *p < 0.05, **p < 0.005.

### Whole genome comparison reveals four *C. innocuum* clades

We next sought to identify genomic heterogeneity among our *C. innocuum* isolates (n = 38) and *C. innocuum* genomes available on GTDB (n = 71), which included genomes sequenced in previously published studies (41, 68) and four fully annotated genomes (*C. innocuum* 14501, I46, LC-LUMC and 2959) (87–89) (Table S1). Isolates were sequenced using Illumina technology, assembled (average depth of coverage = 90X), and annotated using Prokka (67). Five genomes were removed as they were 100% identical (Table S2), leaving 108 *C. innocuum* genomes for further analysis. We compared the newly assembled full-length 16S rRNA gene using both NCBI BLAST and EZBiocloud, and the full genomic assembly against the GTDB database, confirming that all isolates were classified as Erysipelotrichiaceae species. An initial maximum likelihood tree of the full-length 16S rRNA gene obtained from the whole genome assemblies revealed that most species clustered under one branch even when compared to two *C. difficile* 16S rRNA sequences as an outgroup (Figure S1A), suggesting that the 16S rRNA gene may not be an appropriate proxy for distinguishing *C. innocuum* strains.

We next analyzed the pangenome from *de novo* assembled whole genomes of all putative *C. innocuum* genomes available (n = 108) using Roary (95% blast percentage identity) (Table S1) (72). Assemblies of 108 unique genomes averaged 4.6 Mbp, close to the type strain *C. innocuum* ATCC 14501 at 4.7 Mbp (88). We observed an average of 4400 protein encoding genes by coding sequences (CDS) obtained from Prokka (67). The gene accumulation curve followed Heap’s Law with γ = 0.308278 ± 0.05 (R^2^ = 97%) (Figure 2A). Heap’s decay parameter, alpha, totaled less than one (alpha = 0.795), suggesting an open pangenome for *C. innocuum*. This was further supported by the distribution of gene abundance across the number of genomes (Figure 2B), which demonstrated that the number of unique genes (561 genes) common to all 108 genomes was less than those observed in a single genome, indicating extensive gene transference within and outside the species.

**Figure 2.**
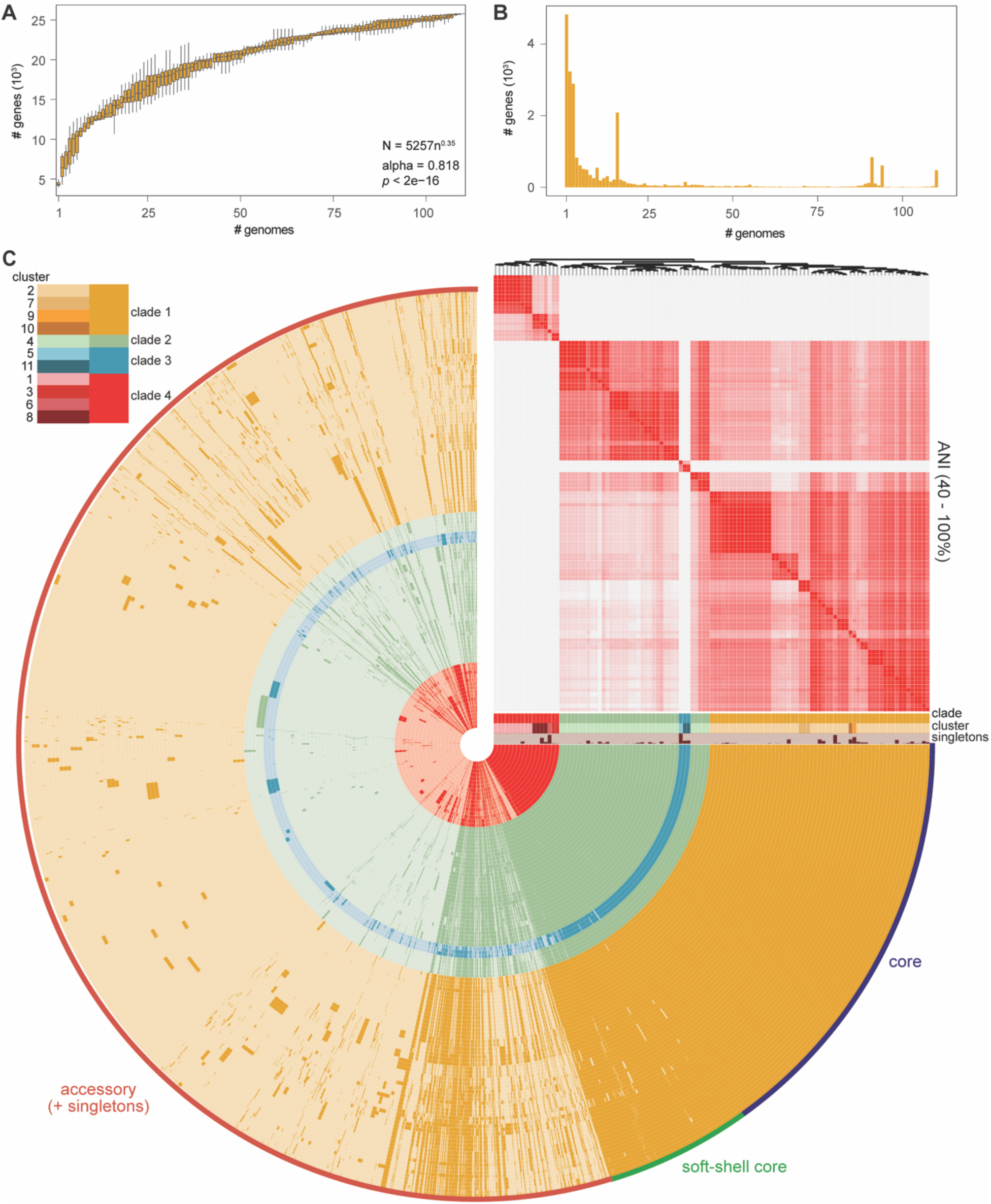
*C. innocuum* exhibits an open pangenome. **A)** Mean number of genes (10^3^) as a function of total number of genomes from five subsamplings (N = 108; alpha = Heap’s law estimate over 500 iterations, estimating the decay parameter fitting the Heap’s Law model (83), p-value < 2e^-16). **B)** Total number of unique genes (10^3^) as a function of number of genomes. **C)** *C. innocuum* pangenome displaying presence/absence of core (purple; present in 100% of genomes), soft-shell core (green; present in 95-99% of genomes), and accessory (red; present in <95% genomes) genes, with hierarchical clustering based on average nucleotide identity (ANI) in right-hand corner (coloring based on 40-100% similarity). Clade and cluster designated in the legend. Analyses incorporate all available unique *C. innocuum* genomes (n = 108).

To investigate genetic variability, we computed the average nucleotide identity (ANI) (57), revealing four distinct clades. Clades III and IV were 90% or less similar to clades I and II, which is less than an expected cut-off for a genus- (90%) or species-level (95%) comparison (90, 91) (Figure 2C). This suggests that clade IV genomes may be associated with a neighboring Erysipelotrichial species instead of the canonical *C. innocuum* species, despite high similarity (99-100%) of the full length 16S rRNA gene alone. Clades I and II were more closely related with >95% ANI (Table S2) and included all four available reference genomes. We then used Anvi’o to assign core (present in all), soft-shell (present in 95 - 99% of genomes), and accessory genes (present in less than 95% of the genomes), which totaled 561 core and 25,176 accessory genes across all 108 genomes (Figure 2C). When excluding clade III and IV genomes, these totaled 2265 core and 14,676 accessory genes, with the pangenome still open as indicated by the alpha and γ parameters (Figure S2).

A maximum likelihood tree using the core genome of all strains reiterated the results observed by ANI (Figure 3A). Unlike the 16S rRNA-based tree, core genome comparison revealed genomic heterogeneity across all available genomes, clustering the strains once again into four major clades (based on 100% bootstrapping for clades I, II, and III, and 65% for clade IV using 500 possible trees). A maximum likelihood tree based on a set of 400 selected protein markers present across all bacterial and archaebacterial constructed using PhyloPhlAn (74, 92) maintained the overall integrity of the four clades, with some shuffling between the closely related clades I and II (Figure S1B).

**Figure 3.**
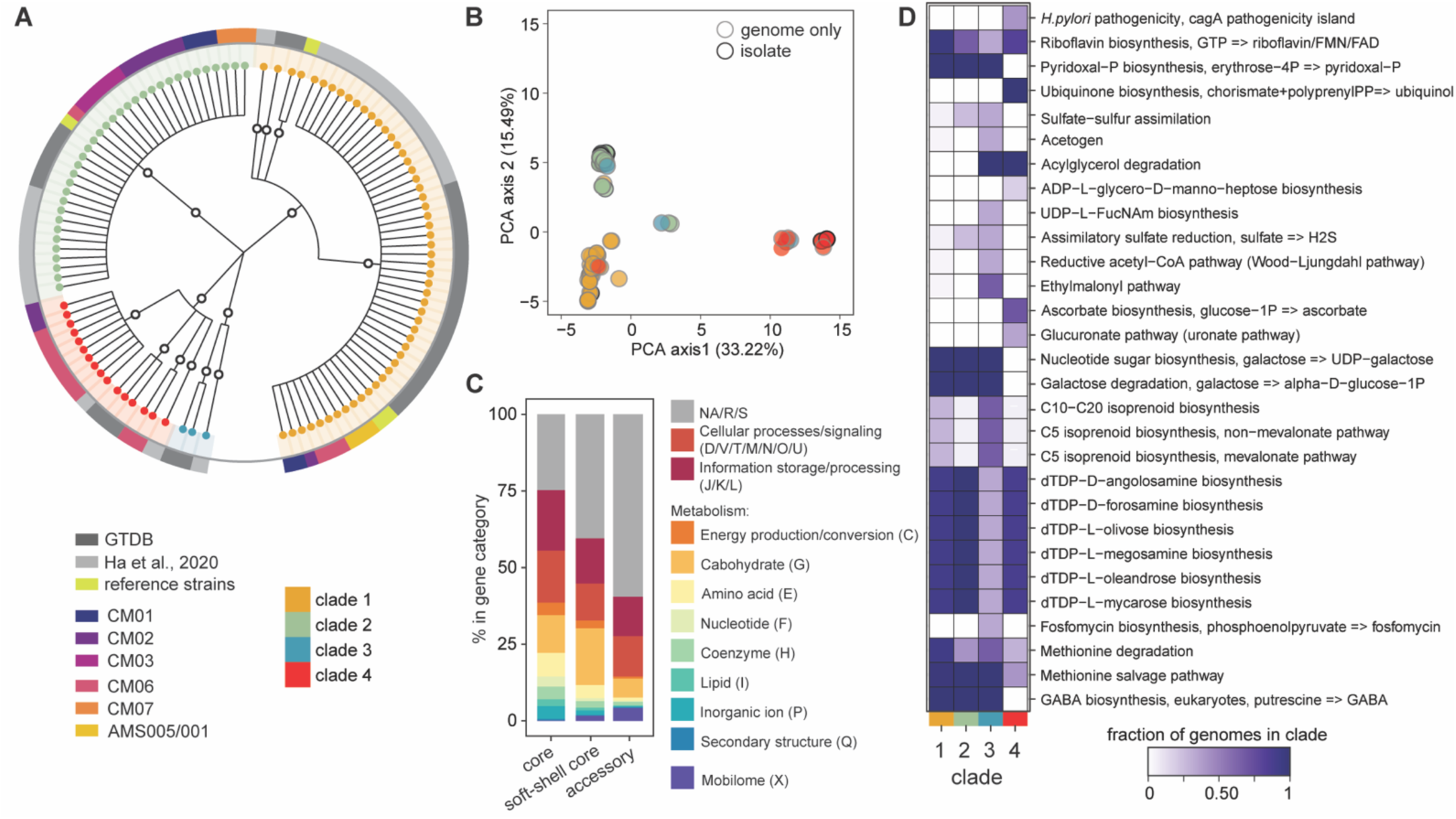
Clade-specific differences in metabolism and potential virulence drive genomic differences in *C. innocuum* strains. **A)** Maximum likelihood tree based on single nucleotide polymorphisms in 400 core genes from 500 replications, colored by clade (node color) and source (circle color) as specified in the legend. **B)** Principal components analysis (PCA) based on covariance from presence/absence of COG gene assignments generated using Prokka colored by clade (legend). **C)** Relative abundance of major COG categories (color-coded in legend) represented in core, soft-shell core, and accessory genes. **D)** Differentially enriched KEGG modules across clades (False Detection Rate using an adjusted q-value below 0.05), colored by fraction of genomes within each clade.

### Genomic differences within *C. innocuum* and related strains is driven by metabolism

Principal Components Analysis (PCA) based on the presence/absence of COG genes demonstrated clustering of strains by clade, with higher similarity of clade III strains to clades II and IV (Figure 3B). Overall, the pangenome of *C. innocuum* displayed ∼25,000 gene clusters categorized using the COG database (Figure 3C). At least 20% of the core genome and 50% of the accessory genes were classified as general functions (R), unknown functions (S) or did not map to the COG database (NA). Genes responsible for basic cellular processes and information storage and processing were equally distributed between the core, soft-shell core, and accessory genomes. Genes involved in metabolism (C, E, F, H, and P) were highest in the core genome, and included genes associated with basic energy producing pathways like ATP synthesis, gluconeogenesis, TCA cycle, urea cycle, or Entner-Duodonoff pathway. From these functions, carbohydrate metabolism (G) held the highest percentage across core, soft-shell core and accessory genes.

Interestingly, genes in the mobilome COG category (X) were over-represented in the accessory genome of *C. innocuum*. Core mobilome representation included only a single mobilome gene, bacteriophage protein gp37, which forms a fibrous parallel homotrimer at the end of the long tail fibers in bacteriophages (93). Some phage-related genes, such as holin (lysis protein), killer protein of prophage maintenance systems (Doc), predicted transposases (InsQ, InsG, and Tra8), and competence proteins (ComCG) were present in the soft-core genome. The majority of the mobilome genes were present in the accessory genome, comprising a multitude of predicted transposases and phage-related regulatory proteins. This included a plasmid stabilization system protein ParE identified across clades I, II and IV strains, which in *Enterobacteriaceae* confers heat and antibiotic tolerance by maintaining IncI and IncF-type plasmid (94).

To identify completeness of metabolic pathways (>75% of total genes), we used Anvi’o for metabolic reconstruction of the strains using the KEGG database. Completed carbohydrate metabolism pathways across all genomes (Figure S3) included the pentose phosphate pathway, pyruvate oxidation, Embden-Meyerhof pathway and Entner-Duodroff pathway. All four clades demonstrated incomplete TCA cycle pathways. Several completed amino acid metabolism pathways were identified across all clades (including valine, proline, and ornithine) except for clade I, which lacked a complete module for cysteine. For lipid metabolism, completed pathways included fatty acid biosynthesis, initiation, and elongation.

We used the Anvi’o functional enrichment tool to identify differentially abundant KEGG modules across clades (p-value < 0.05). Overall, clade I and II genomes exhibited similar profiles to each other compared to clade III and IV, although clade III and IV genomes also exhibited differences. (Figure 3D). Clades I and II shared genes related to biosynthesis of terpenoids and polyketides that were less represented in clades III and IV. The glucuronate (uronate), ascorbate biosynthesis, ubiquinone biosynthesis and ADP-L-glycero-D-manno-heptose biosynthesis pathways were almost exclusively represented within clade IV strains compared to other clades. These pathways consisted of only two KOfam assignments, UDP-glucose-6-dehydrogenase [E.C. 1.1.1.22] and xylulokinase [E.C. 2.7.1.17], suggesting incomplete pathways for biosynthesis of ascorbate and the pentose phosphate metabolic pathway, respectively.

Clinically relevant clade differences were also identified. While module representation across all strains suggest that lipid biosynthesis and utilization is prevalent within *C. innocuum* (Figure S3), acylglycerol degradation (triacylglycerol lipase, [E.C.: 3.1.1.3]) was observed for only strains in clade III and IV. Our analysis included *C. innocuum* genomes (SRR12535151 and SRR12535143) recently isolated from creeping fat in patients with Crohn’s disease as belonging to clade IV (41). A *Helicobacter pylori cagA* pathogenicity island signature module (K02283) was also identified as differentially represented in clade IV genomes. This includes the type IV pilus assembly protein, CpaF [E.C: 7.4.2.8].

To identify more specialized polysaccharide metabolism in *C. innocuum*, we used dbCAN and the CAZy database to identify carbohydrate-active-enzymes (CAZymes) (84). Glycoside hydrolases in the GH1 group and GT2 glycosyl transferases were the most abundant CAZymes in *C. innocuum* collectively (Figure 4). Strains in clades I and II encompassed the most diverse and abundant CAZyme compared to the other two clades. Compared to clade II strains, strains in clade I were more likely to contain multiple glycoside hydrolases involved in alpha- or beta-D-glycosidic bond hydrolysis (GH2, GH43_35, GH65, GH106, GH140, and GH146). Compared to either clade I or II strains, strains in clades III and IV demonstrated fewer CAZymes. GT14, which produces a glycogen branching protein, was present in both clades I and IV, while GT11, a fucosyltransferase, was only present in clade IV. The acetyl-mannosamine transferase GT26 was only present in clade II strains. Only three types of carbohydrate esterases (CE) were identified across all *C. innocuum* strains, and only one group of carbohydrate binding modules (CBM66), part of the LPXTG cell wall anchor domain-containing protein, was identified in clades I and IV only.

**Figure 4.**
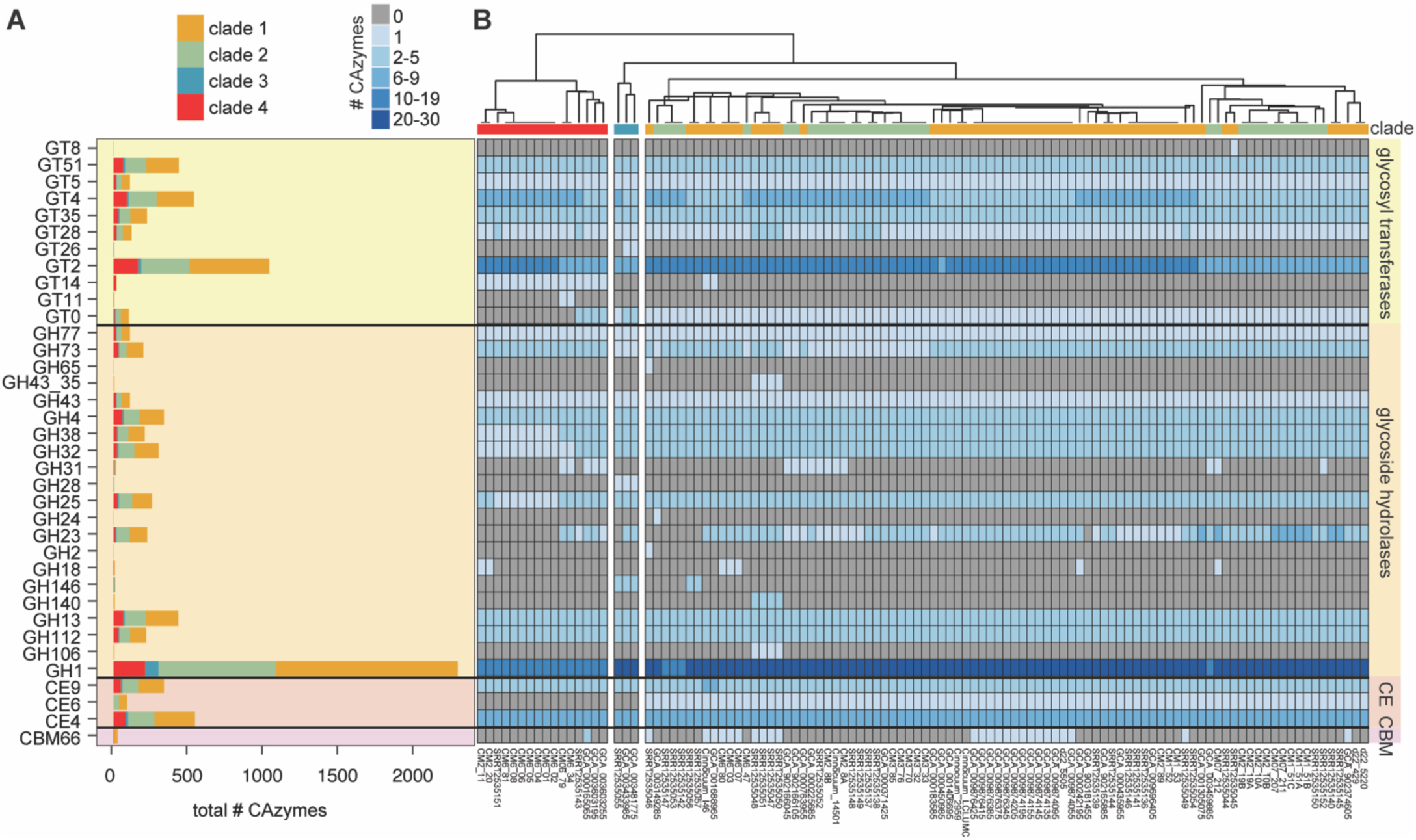
*C. innocuum* strains display clade-specific carbohydrate-activating enzymes (CAZymes). Number and type of CAZymes detected in *C. innocuum* genomes, color-coded by clade type. **B)** Heatmap of number and type of CAZymes in individual genomes, clustered using Euclidean distance. GT = glycosyltransferase, GH = glycoside hydrolase, CE = carbohydrate esterase, CBM = carbohydrate-binding module.

**Figure 5.**
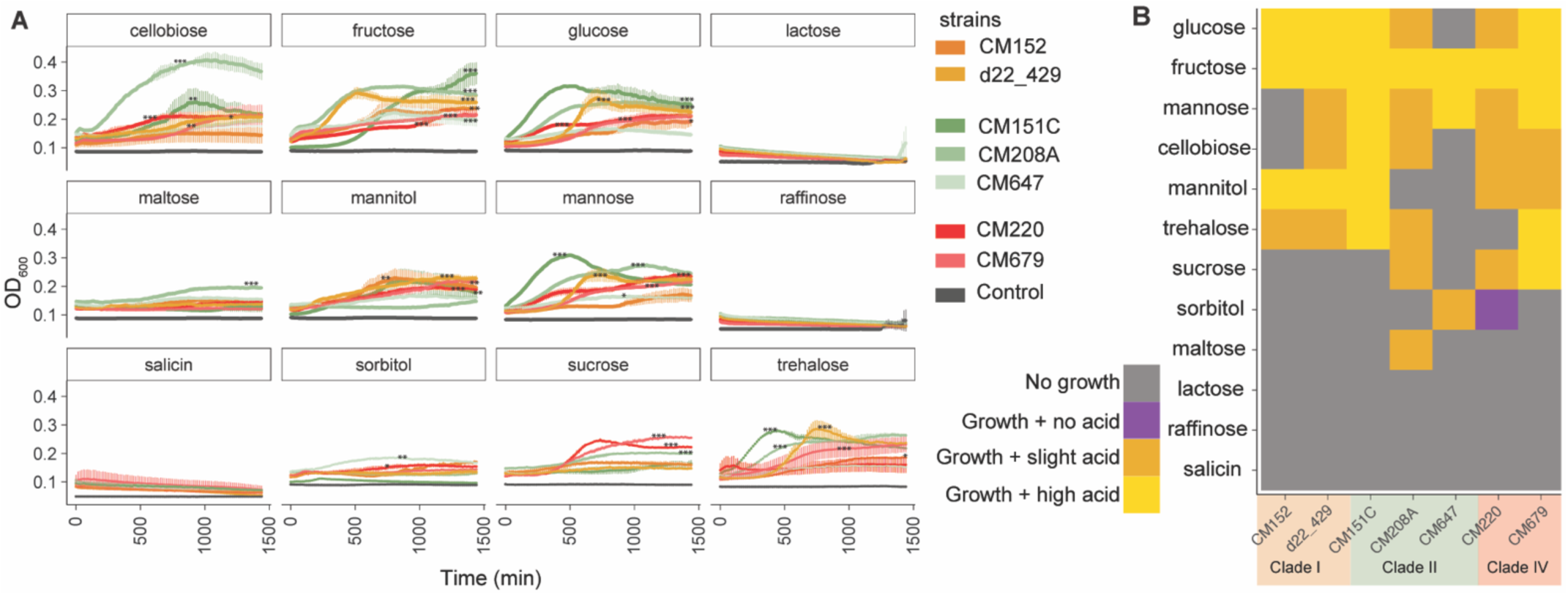
*C. innocuum* exhibits strain-specific differences nutrient utilization *in vitro*. **A)** Growth curves of strains (n = 7) inoculated into basal media with casamino acids (BMCA) with indicated carbohydrate source over 24 hours, measured at OD_600_. Grey indicates the negative control (strain inoculated into BMCA without addition of carbohydrate). Significance of growth was determined by ANOVA on area under curve per strain (within each sugar type), followed by Tukey’s HSD, *, p < 0.05; **, p < 0.005; ***, p < 0.0001. **B)** Growth (from OD_600_) and acid production (from bromocresol purple assay) data combined to show nutrient utilization patterns in representative strains. High acid production was classified as pH < 5.5 and OD < 0.26, low acid production was classified as 5.5-6.5 and OD_588_ 0.26-0.44 (legend). Clade/strain designation indicated by legend.

### *C. innocuum* exhibits strain-level variation in substrate use *in vitro*

To examine differences in nutrient use across strains, we selected seven representative strains to examine their ability to use distinct carbohydrate sources *in vitro*. Representative strains were selected from a 99% dereplication cutoff of *C. innocuum* genomes, which clustered the genomes into seven groups, representing three of the four clades (Table S2). We assessed both growth and acid production of the strains in minimal media supplemented with single carbohydrates (Table S3). After 24 hours of growth assessment, acid production was assessed using the colorimetric pH indicator, bromocresol purple (BCP), which approximates pH changes as a result of fermentation (Figure S4). Acid production signifying fermentation was determined as low (5.5-6.5 and OD_588_ 0.26-0.44) or high (< 5.5 and OD_588_ < 0.26).

We observed variation in the ability of strains to grow on minimal media supplemented with single carbohydrate sources (Figure 7). None of the strains were capable of growing on lactose or raffinose, confirming previous observations for *C. innocuum* (28, 96). In contrast to previous reports (33), salicin did not support growth of any of the strains tested. Most strains demonstrated high growth (defined by both significant increases in OD and high acid production) in glucose and fructose, albeit at varying growth rates. Significant growth on glucose and mannose was observed by all but one strain, CM6_47 or CM1_52, respectively (ANOVA on AUC, p < 0.05). Cellobiose, mannitol and trehalose all supported growth and acid production in a majority of strains (ANOVA, p < 0.05). Maltose supported the growth of only one strain, CM208A (ANOVA, p < 0.05), whereas sorbitol only supported growth of two strains, CM647 and CM220 (ANOVA p < 0.05).

Only some of the variable growth aligned with their clade designation. Within clade I, both CM1_52 and d22_429 followed a similar pattern of acid production (Figure 7B) and grew at varying efficiencies in glucose, fructose, mannitol, and trehalose (Figure 7A). Within clade IV, both strains CM2_20 and CM6_79 grew efficiently in several carbohydrates, and more efficiently on sucrose than most strains (ANOVA, p < 0.0001). The most variation was observed in clade II strains, with CM6_47 consistently displaying minimal growth on most carbohydrates. In contrast, both CM2_08A and CM1_51C exhibited some of the highest growth in glucose, cellobiose, and mannose compared to other strains (ANOVA, p < 0.005), but CM2_08A did not grow in mannitol compared to CM1_51C (ANOVA, p < 0.005). CM2_08A also consistently produced less acid than CM1_51C with the exception of growth in fructose (Figure 7B).

### Virulence and antibiotic resistance genes in *C. innocuum* are clade-dependent

An exotoxin has not been identified from *C. innocuum* despite previous evidence association with infection (97). Using PathoFact to identify potential virulence (bit score > 50), we observed differential distribution of putative virulence factors across the four clades (Figure 6A) (85). Overall, clade III strains had lower numbers of virulence factors detected over the other clades, which also lacked type II toxin antitoxin system from the YafQ/RelB/ParE family and phage lysis holins. Some factors were present across all clades, including members of hemolysin III, *hlyIII* and *tlyC*; NlpC/P60 family, *pspA* and *pspC* (identified as *entD* in the software); a type III toxin-antitoxin system from ToxN/AbiQ (Figure 6A). GGGtGRt, *tlyC*, *hlyIII* demonstrated the highest bit scores across all clades, including within the reference strain *C. innocuum* 14501. While we identified the presence of *C. difficile tcdAB*-like genes in clades I and II, BLAST search across the NCBI databases revealed that they likely belong to the NlpC/P60 family of proteins, either as surface proteins *pspAC* alongside a penicillin binding protein *mrcB* or as a glucan-binding domain containing protein not yet fully characterized.

**Figure 6.**
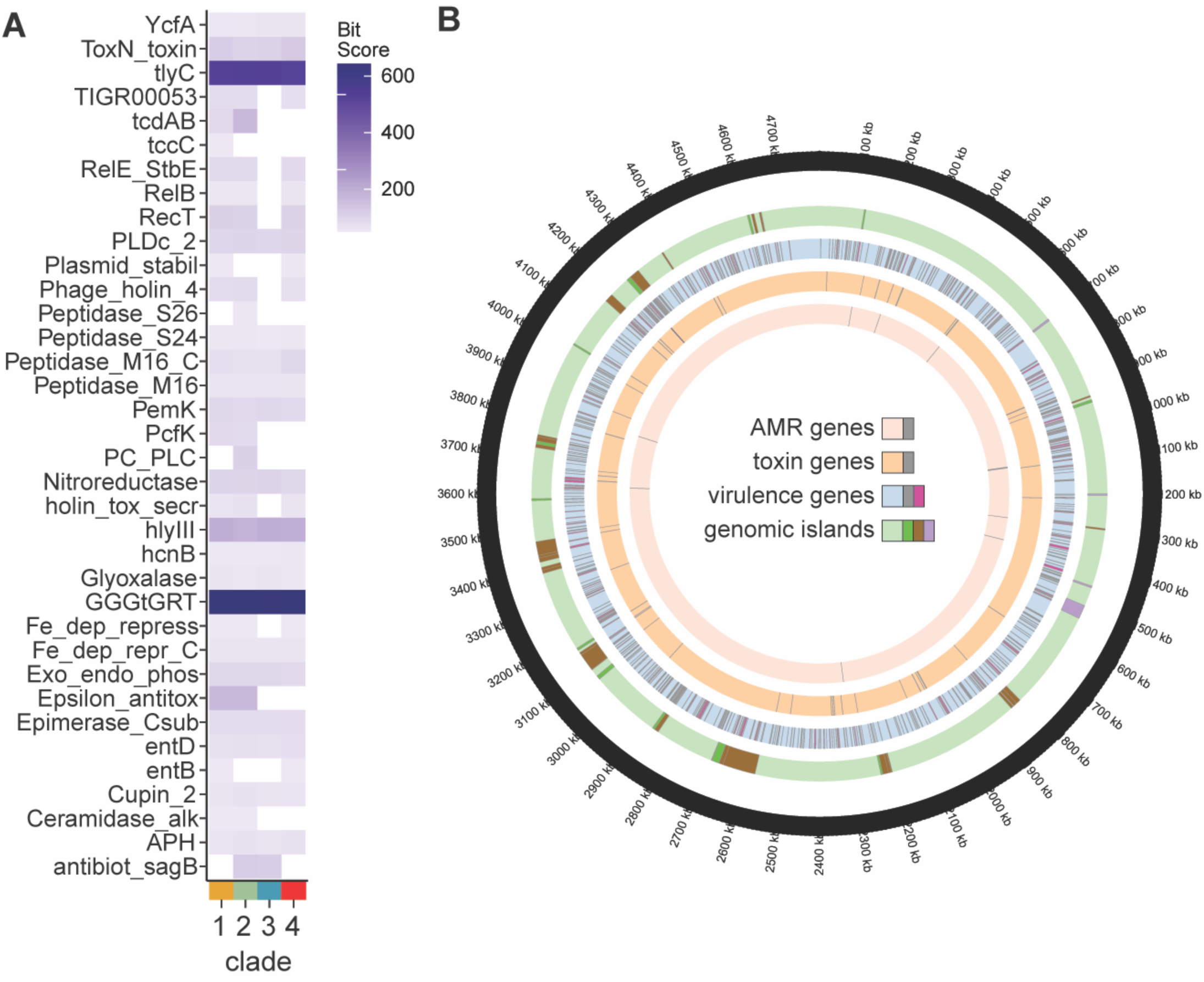
Certain clades of *C. innocuum* display enhanced potential virulence. **A)** Average bit score of select putative toxins (bit score > 40) across *C. innocuum* clade, identified using PathoFact. **B)** Location of antibiotic resistance genes (AMR), putative virulence factors and toxins identified using PathoFact (three inner-most circles) and genomic islands, identified using IslandViewer4 (penultimate outer circle; green = forward orientation; brown = reverse; purple = bidirectional orientation).

The chromosomal location of virulence and antibiotic resistance genes identified by PathoFact were visualized in conjunction with genomic islands (using Island-Viewer4), with *C. innocuum* 14501 as a reference (Figure 6B) (86). Resistance genes against vancomycin (*vanRS)*, tetracycline, as well as several ABC transporters and aminoglycoside genes (Table S5) were identified in all clades. Bacitracin resistance (*bcrAC*) was found only in clades I and III (Figure S5). Results from Island-Viewer4 predicted 42 genomic islands, with 113 virulence factors and seven toxin genes aligning with genomic island locations (Figure 6B). While none of the hemolysins aligned with genomic island predictions, a type II TA system involving RelE/StbE family of toxin-antitoxin system (with 11 additional genes) and a group of NlpC/P60 glucan binding proteins (labeled as *tcdAB* by PathoFact, with 9 additional genes) each aligned with a predicted genomic island.

## DISCUSSION

To date, this study marks the most comprehensive characterization of the human gut inhabitant, *C. innocuum*. While *C. innocuum* has originally been designated a commensal from initial isolation studies (33, 35), it has also been recently associated with various diseases states (30, 38–41). We recovered *C. innocuum* strains from all individuals sampled in this study, suggesting higher prevalence of *C. innocuum* in the ‘healthy’ human gut than previously expected. This is further strengthened by the prevalence of *C. innocuum* in human 16S rRNA gene-based surveys using a custom classifier specific for *C. innocuum* 16S rRNA sequences. *C. innocuum* is rarely identified in studies using human 16S rRNA surveys, either in association with health or disease. This may be due to its low abundance in comparison to other microbes or misclassification of the short 16S rRNA gene fragment using standard databases. Nevertheless, the consistent association of *C. innocuum* with disease in culture-based studies, as well as the observed increased abundance of *C. innocuum* following antibiotic use in our specific 16S rRNA survey, support a closer look at the role of *C. innocuum* in the gut microbiota.

Our genomic analysis of *C. innocuum* clarifies some of the functions attributable to *C. innocuum* colonization of the gut. We identified several complete modules in both carbohydrate and amino acid metabolism, such as pentose phosphate pathway, pyruvate oxidation, Embden-Meyerhof pathway and Entner-Duodroff pathway, and various amino acid biosynthesis and metabolism pathways in the *C. innocuum* core genome. These also included multiple genes associated with utilization of saccharides such as glucose, mannose, fructose, xylose, and mannitol, indicating an ability to use byproducts of polysaccharide degradation by other commensals. Compared to other anaerobic bacteria commonly isolated alongside *C. innocuum* such as *C. difficile*, *C. acetobutylicum*, and *C. kluyveri* (99), *C. innocuum* displayed an incomplete TCA cycle. Additionally, all *C. innocuum* strains exhibited several partial and complete modules for lipid metabolism. None of the tested strains were able to grow in lactose, salicin and raffinose, corroborating descriptions of *C. innocuum* clinical isolates growth using a Biolog platform (41, 100).

Our genomic comparison also reveals strain-specific diversity in *C. innocuum*. After dereplicating *C. innocuum* strains to remove genetically identical strains from our dataset, both ANI and phylogenetic analysis of *C. innocuum* genomes demonstrated four distinct clusters. Clades III and IV were collectively 90% or less similar to clades I and II, which contained all available *C. innocuum* reference strains, suggesting that this clade may represent a new Erysipelotrichial species related to *C. innocuum*. Even after exclusion of clade III and IV genomes, the pangenome of *C. innocuum* remains highly open as assessed by Heap’s law (83). It has been suggested that an open genome may be more reflective of a sympatric lifestyle, whereby related species interacting with each other can easily share genetic elements (101). While our current study did not specifically focus on identification of mobile elements, most mobilome-related genes were present in the accessory portion of the *C. innocuum* pangenome, suggesting a high degree of horizontal gene transfer among *C. innocuum* and related species.

The ability to acquire new genes can provide a competitive metabolic advantage in a microbially dense environment. As the nutrient niche theory stipulates, colonization by an invading microbe, pathogenic or commensal, is at least partially dependent on its ability to outcompete extant microbes in that environment (102). For example, co-existence of the highly abundant gut inhabitant *Bacteroides thetaiotaomicron* is likely possible at least in part due to the diversity of polysaccharide-utilizing loci observed across different strains that allow flexibility in resource utilization (103). We observed co-existence of several *C. innocuum* strains within a single fecal sample, none of which were 100% identical to each other and some of which spanned multiple clades within an individual. Overall, the CAZymes observed across *C. innocuum* were fewer in number compared to previously characterized gut commensals (14, 104, 105). However, we did observe distinct clade-specific clustering based on CAZyme presence, with clades I and II demonstrating increased CAZyme diversity and abundance compared to clades III and IV. These genome-encoded CAZyme differences between clades and potential new species could indicate niche partitioning to support related strains within the same environment.

The realized, or expressed, niche of strain co-existence is likely more complicated (106). Our *in vitro* growth assays support niche partitioning within the canonical *C. innocuum* clades I and II at both clade- and strain-specific levels. For example, both CM01-52 (clade I) and CM01-51C (clade II), isolated from the same individual, could use fructose, glucose, mannose, and trehalose for growth. Yet, the former grew significantly better with fructose, whereas the latter grew better on glucose, mannose, and trehalose. Despite the observation of clade-specific genomic patterns in CAZymes, we did not observe overt clade-specific growth patterns *in vitro*. This suggests that realized metabolic niche partitioning can occur within an individual beyond the categorical genomic features assessed, emphasizing the importance of regulatory or additional genes that contribute to successful co-existence of related strains. These differences also likely influence or are influenced by other members of the collective microbiota within an individual.

The demonstrated genomic and phenotypic variability observed across the *C. innocuum* species may also be of clinical importance. *C. innocuum* is commonly isolated in conjunction with gastrointestinal tissue or fecal clinical samples (36, 41, 97). A recent retrospective study in a Taiwanese clinical cohort isolated *C. innocuum* rather than *C. difficile* from patients with *C. difficile*-like clinical presentation (36). These *C. innocuum* isolates were reported to cause a range of cytotoxicity and enteropathogenic effects *in vitro*. Most recently, Cherny et al. reported that *C. innocuum* isolates from pediatric patients enrolled in *C. difficile* studies cross-reacted with the enzyme immunoassay (EIA) diagnostic test for *C. difficile* toxins A/B. The study identified a putative *C. innocuum* toxin EIA cross-reactivefactor (ErF) similar to the NlpC/P60 family of toxins in all isolates tested but observed no cytotoxicity. We identified the same putative toxin A/B gene in clades I and II, with a significantly higher similarity score for tcdA/B in clade II genomes. *C. innocuum* has also been postulated as a potential causative agent of ‘creeping fat’, an extra-intestinal phenomenon correlated with Crohn’s Disease (CD) (41). Ha et al. demonstrated that *C. innocuum* isolated from various human intestinal mucosal locations could translocate into tissue in a mouse model of IBD. Our analysis, which included genomes from that study, did not identify clade-specific clustering based on the anatomical site. However, the two clade IV representatives identified as part of this study were both isolated from creeping fat lesions. Together, these data suggest the possibility of strain-specific *C. innocuum* virulence that may explain ambiguity in identifying *C. innocuum* or a closely related species as an opportunistic or absolute pathogen.

In summary, we demonstrate strain-specific variation of a prevalent gut ‘commensal’ that until recently was considered relatively benign in the gut environment. The increased association of *C. innocuum* with gastrointestinal conditions support further investigation of specific virulence features that may contribute to human disease, with an emphasis on explicit identification of potentially pathogenic or commensal *C. innocuum* strains. Furthermore, our results reveal the importance of understanding strain variation that can be extended to other gut commensals.

## ACKNOWLEDGEMENTS

We would like to thank the donors who consented to donating their fecal samples for our study. We would like to acknowledge Clemson University for generous allotment of compute time on Palmetto cluster. This publication was made possible, in part, with support from the Clemson University Genomics and Bioinformatics Facility, which receives support from an Institutional Development Award (IDeA) from the National Institute of General Medical Sciences of the National Institutes of Health under grant number P20GM109094. AMS was supported by grant number K01-DK111794 from the National Institute of Diabetes and Digestive and Kidney Diseases. We would like to acknowledge Microbial Genome Sequencing Center (MiGS) for whole genome sequencing of our *C. innocuum* strains, and Eton Biosciences for Sanger sequencing the amplified 16S rRNA genes. AMS has received consultation fees from Finch Therapeutics and Rebiotix / Ferring Pharmaceuticals.

D.B. - Conceptualization, Data Curation, Formal analysis, Investigation, Methodology, Software, Writing—original draft, and Writing—review and editing; C.F. – Data Curation, Formal analysis, Investigation, Methodology, Validation, Writing – original draft; C.W.M – Methodology, Investigation, Writing – original draft; A.M.S. – Supervision, Project Administration, Funding Acquisition, Conceptualization, Writing – original draft, Writing – review and editing.

